# Habitat use and diel activity of insectivorous bats across land-use types on an Afrotropical oceanic island

**DOI:** 10.1101/2025.04.10.648189

**Authors:** Ana Filipa Palmeirim, Ana Catarina Araújo-Fernandes, Ana Sofia Castro-Fernandes, Patrícia Guedes, Yodiney dos Santos, João C. Alves, Vanessa Mata, Natalie Yoh, Ricardo Rocha

**Affiliations:** CIBIO, Centro de Investigação em Biodiversidade e Recursos Genéticos, InBIO Laboratório Associado, Campus de Vairão, Universidade do Porto, 4485-661 Vairão, Portugal; BIOPOLIS Program in Genomics, Biodiversity and Land Planning, CIBIO, Campus de Vairão, 4485-661 Vairão, Portugal; EcoHealth Alliance, New York City, NY, United States; Departamento de Biologia, Faculdade de Ciências, Universidade do Porto, 4169-007, Rua Campo Alegre s/n, Porto, Portugal; Fundação Príncipe, Santo António, Ilha do Príncipe, São Tomé e Príncipe; Durrell Institute of Conservation and Ecology (DICE), School of Anthropology and Conservation, Canterbury, UK; Department of Biology, University of Oxford, 11a Mansfield Rd, OX1 3SZ Oxford, UK

**Keywords:** human-modified landscapes, Chiroptera, endemic species, Gulf of Guinea, land-use change, tropical forests, passive acoustic monitoring, São Tomé, Príncipe

## Abstract

Tropical island biodiversity is declining at alarming rates. Yet, understanding how species are coping with such disturbance is largely limited for afro-tropical islands. Here we examined habitat use and diel activity of insectivorous bats across different land-use types covering the endemic-rich Príncipe Island, Central West Africa. We acoustically surveyed insectivorous bats across 48 sites throughout old-growth forests, secondary re-growth forests, cocoa shaded plantations, and horticultures. Based on 17,527 bat-passes, we were able to record all four insectivorous bat species known to occur on Príncipe, including the recently described *Pseudoromicia principis*, the most frequently recorded species. *Taphozous mauritianus*, a data deficient open-space forager, was the least recorded species. Species activity was lowest in old-growth forests, as well as the activity of the edge-forager *P. principis*. The activity of the open-space forager *Mops pumilus* was lowest in forest habitats, whereas *Hipposideros ruber*, a forest forager, was restricted to these habitats. Diel activity patterns of *M. pumilus* varied between forests and non-forest habitats, whereas those of *P. principis* remained similar. Interspecific activity overlap decreased towards more altered land-use types. Our findings emphasize that conserving the remaining forests, along with the current mosaic of land-uses, is needed to maintain Príncipe’s complete insectivorous bat assemblages.

## 1. INTRODUCTION

Land-use change is the primary driver of biodiversity loss worldwide (Caro et al., 2022). This is especially concerning in the humid tropics, where deforestation— primarily due to agricultural expansion—continues to erode vast tracts of native habitat (Hansen et al., 2020). The situation is even more critical on tropical oceanic islands, as their often-high levels of endemism (Fernández-Palacios et al., 2021), greater physical and behavioural specialization among species (Kier et al., 2009), and heightened exposure to natural hazards (Gonçalves et al., 2024) makes their unique biota particularly vulnerable to habitat disturbance.

The effects of land-use change on biodiversity can be complex, depending on both the intrinsic species traits (Newbold et al., 2016) and the degree of structural and compositional change in relation to the baseline habitat (Barlow et al., 2007). Following anthropogenic disturbance, while some disturbance-sensitive species might undergo extinction (Palmeirim et al. 2017), the persisting species can adjust their habitat use and behaviour in the novel environmental conditions (Mazza et al., 2020; Tranquillo et al., 2023). More profoundly altered habitats also tend to have more contrasting conditions, including variations in resource availability, microclimate, light duration (e.g., increased exposure in newly open areas), human presence, shifts in species composition (Fletcher et al., 2024), and greater inter-and intraspecific competition (Monterroso et al., 2014). These factors might synergistically act to promote species changes in habitat use (Ferreira et al., 2022) as well as changes their diel activity patterns (Ikeda et al., 2016). Understanding how species cope with land-use change remains a cornerstone to outline evidence-based management actions, which is not trivial given the accelerated pace of land-use change across tropical islands (Wood et al., 2017).

On oceanic islands, bats often comprise the only native mammal species (Fleming & Racey, 2009), with ca. 25% of all bat species occurring on islands being island-endemics (Conenna et al., 2017). Despite the high conservation concern of insular bats, bat responses to land-use change have rarely been explored on afrotropical islands (but see e.g., Kemp et al., 2019 and Mandl et al., 2021). Overall, bat responses to land-use change are typically driven by features of their echolocation call characteristics, diet, and morphology (Davies et al., 2016; Wordley et al., 2017; Núñez et al., 2019). In aerial insectivorous bats, echolocation characteristics are intrinsically linked to their habitat affinities and foraging strategies, and according to those, aerial insectivorous bats are often categorized according to three main foraging guilds: forest, edge and open-space foragers (Schnitzler & Kalko, 2001; Yoh et al., 2022). Forest foragers emitting high-frequency constant frequency (CF) or frequency-modulated (FM) echolocation calls generally comprise the most vulnerable species to land-use changes that imply a decline in the habitat clutterness (Hazard et al., 2023; Rowley et al., 2024). On the other hand, edge and open-space forager bats typically use frequency-modulated, quasi-constant frequency calls. As such, edge forager bats might benefit from the creation of edges while open-area forager bats might benefit from the removal of the tridimensionality habitat complexity (Mendes & Srbek□Araujo, 2021; López□Bosch et al., 2021).

In the Atlantic coast of Central Africa, Príncipe Island (139 km^2^) is part of the Cameroon volcanic line in the so-called Gulf of Guinea (Ceríaco et al., 2022a). Despite the relatively short distance from mainland Africa (220 km), it remained isolated throughout its geological history possibly starting approximately 31 Ma years ago (Ceríaco et al., 2022b). As a result, Príncipe is characterised by exceptional levels of endemism, including eight endemic bird species (29% of the native species), three of amphibians (100%), eight of reptiles (67%) and two of mammals (29%) (Ceríaco et al., 2022b). The island was permanently colonised by humans since the 15^th^ century and ca. 25% of its area remains covered by native old-growth forests (Dauby et al., 2022). The rest of the island consists of a mosaic of land-use types, dominated by abandoned plantations that have regenerated into secondary re-growth forests, shaded cocoa plantations, small horticulture holdings, and urban areas (Dauby et al., 2022). Nonetheless, despite the island’s relevance for conservation, and its high degree of anthropogenic influence, the impacts of land-use change on Príncipe’s native fauna remains largely unexplored (but see Dallimer & King, 2007 and Rebelo et al., 2024).

In this study, we examine how insectivorous bats utilise the various land-use types across the endemic-rich Príncipe Island. The island’s insectivorous bat assemblage is composed by four species: *Hipposideros ruber* (a Hipposideridae and forest forager), *Pseudoromicia principis* (an endemic Vespertilionidae and edge forager recently described by Juste et al., 2023), *Mops pumilus* (a Molossidae and open-space forager), and *Taphozous mauritianus* (an Emballonuridae and open-space forager) (Rainho et al. 2022). Specifically, we analysed habitat use and diel activity patterns of each species across the island’s four predominant land-use types: old-growth forests, secondary re-growth forests, cocoa shaded plantations, and horticultures. Given the diversity foraging preferences among Príncipe’s insectivorous bats, we expected considerable changes in species composition between land-use types due to species-specific responses. We anticipate *H. ruber* to be mostly associated with forested land-use types, while the remaining species are likely to exhibit higher activity in more altered land-uses such as horticultures, where synanthropic species are likely to benefit from human-associated foraging and roosting resources (Russo & Ancillotto, 2015, Lopez-Baucells et al., 2017). We further expected diel activity patterns to vary most between forested habitats and horticultures, given structural contrast between these habitats, likely resulting in differing light conditions and arthropod diversity (Presley et al., 2009, Montaño-Centellas et al., 2015). Within the same land-use type, overlap in interspecific bat activity is expected to be lower in land-use types where multi-species activity is higher, comprising a temporal partitioning strategy to reduce interspecific competition (Lambert et al., 2018).

## 2. METHODS

### 2.1 Study area

The study was conducted on Príncipe Island, one of the two main islands comprising the Democratic Republic of São Tomé and Príncipe, off the western equatorial coast of Central Africa (Fig. 1). With a human population of ca. 8.000 people, Príncipe Island experiences an average annual temperature of 26 ºC, with minor fluctuations between coastal and mountainous regions (highest point at 948 m a.s.l). The average annual precipitation ranges from 1000 mm in the northeast to around 3000 mm in the south (Ceríaco et al., 2022a).

**Figure 1.**
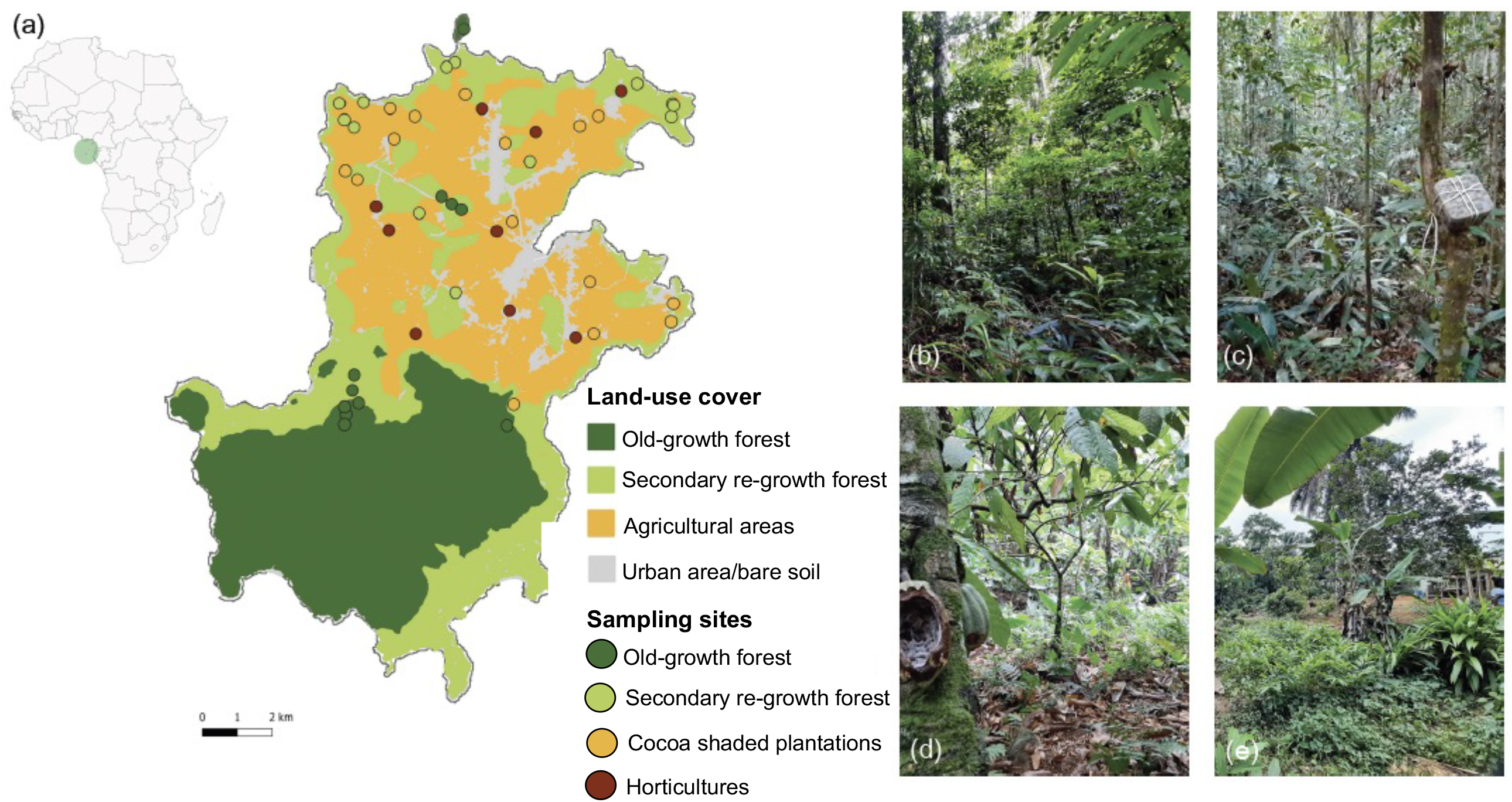
(a) Sampling sites across Príncipe Island, West Central Africa. Each of the 48 sampling sites is colour-coded according to the land-use type: (b) old-growth forest, (c) secondary re-growth forest, (d) cocoa shaded plantation, and (e) horticultures. Land-use types shown in this map were adapted from Soares (2019). Agricultural areas include both cocoa shaded plantations and horticultures.

Príncipe Island is covered by old-growth forests, regrowth forests, shaded plantations, horticultures and, to a lesser extent, urban areas (Soares, 2019). Old-growth forests comprise the least disturbed land-use, covered by native forest vegetation. While old-growth forest may have experienced alterations in the past, these areas are now largely characterised by known human intervention and mostly located within the limits of the Ôbo Natural Park, a significant conservation area covering 65 km^2^ (46% of the island’s area). Secondary re-growth forests largely result from re-growth following the abandonment of cocoa plantations and include the presence of introduced tree species. Part of these forests is also within the limits of the Natural Park. Shaded plantations comprise agroforestry systems where (mostly) cocoa grows under the canopy of predominantly exotic trees. Horticultures refer to small-holding fields growing banana, cassava, carrots, and tomatoes, among others.

### 2.2 Acoustic surveys

Insectivorous bats were acoustically surveyed across 48 sampling sites: 11 sites in old-growth forests, 13 in secondary re-growth forests, 15 in cocoa shaded plantations, and nine in horticultures (Fig. 1). Sampling occurred during August 2022. This period was selected to avoid heavy rain during the surveys, which have the potential to impact both bat activity and the performance of the recorders. The sampling site location aimed to cover as much area of the island as possible. However, due to difficulties in accessing more remote areas, sampling for old-growth forests was geographically restricted (Fig. 1). Sampling sites were at least 250 m apart from each other to minimise any pseudo-replication effect. Each sampling site was surveyed using one AudioMoth acoustic device (Hill et al., 2018), that was attached to a tree trunk at ∼2 m-height. Acoustic devices were set to record 1 min every 5 mins at a sample rate frequency of 384 kHz for 48 h including during the day as *H. ruber* is known to exhibit diurnal flight (Russo et al., 2011).

### 2.3 Bioacoustic analysis

Bat activity was measured using a “bat pass”, representing a sequence of two or more echolocation pulses emitted by a specific species within a five-second audio file (Yoh et al., 2022). The original one-minute recordings obtained from the acoustic devices were divided into five-second files using Kaleidoscope v.5.3.8 software (Wildlife Acoustics, USA). This division allowed for the isolation of files likely containing bat passes, while additional files were automatically sorted into a separate “noise” folder. To select files containing bat passes, specific signal detection parameters were established in Kaleidoscope. This included setting the detection range for frequencies between 8 and 250 kHz, and the minimum and maximum durations of detection pulses between 2 and 500 milliseconds. The software was configured to consider a maximum inter-syllable gap of 500 milliseconds, and a minimum of two pulses were required for detection.

Species were then identified by analysing the characteristics of the echolocation call, namely the call shape, minimum and maximum pulse frequency, peak frequency, call duration, and the interval between pulses. The characteristics of the echolocation call for each species were sourced from Rainho et al. (2010), with specific information regarding *P. principis* obtained from Just et al. (2023). Species were manually identified using Kaleidoscope. Social calls and feeding buzzes were not identified to the species level and thus not considered in this study.

### 2.4 Data analysis

#### 2.4.1 Habitat use

Species activity was examined at the assemblage and species-specific levels, and it was given by the number of bat passes, a proxy of species abundance (Kunz et al., 2009). Species composition across the different land-use types using a Non-Metric Multidimensional Scaling (NMDS) ordination based on a Bray-Curtis similarity matrix, considering the number of bat passes per site for each species (stress: 0.147). We tested whether species composition varied between land-use types using a Permutational Multivariate Analysis of Variance (PERMANOVA). Complementary, we applied a Permutational Analysis of Multivariate Dispersions (PERMDISP) to compare the dispersion in species composition between land-use types. These analyses were carried out using the *vegan* R package (Oksanen et al., 2013).

To examine the effects of land-use type and altitude on overall bat species activity and species-specific activity, we applied Generalised Linear Mixed-effects Models (GLMMs), using the *lme4* R package (Bates et al., 2015). To account for natural variability in bat activity between sampling sites spatially closer and that have been simultaneously sampled, a random term indicating the date the sampling site had been surveyed was used. Altitude was standardised to a mean of zero and a standard deviation of one. The distribution family used in each model varied according to the response variable: Poisson for overall activity and negative binomial for *P. principis, M. pumilus*, and *H. ruber*, while *T. mauritianus* was only detected in three sites and so excluded from subsequent analysis. We further examined the residuals of each model using the *DHARMa* R package (Hartig, 2022).

#### 2.4.2 Diel activity patterns

To assess diel activity patterns, the number of bat passes recorded per species per land-use type was summed for each hour. To ensure robust data representation, only species detected in at least 10% of the surveyed nights with a minimum of 100 independent detections per land-use type (Lashley et al., 2018) were considered in subsequent analysis. As such, we were not able to include *T. mauritianus* nor *H. ruber* in subsequent analysis. The old-growth forests and regrowth forests represented relatively undisturbed areas with similar canopy cover (mean ± SD: 40.83 ± 16.76% in old-growth forests and 25.33 ± 19.13% in secondary re-growth forests at 32-m height; Table S1). Therefore, and due to the low number of bat detections in old-growth forests alone, we decided to merge these two categories of land-use in these and subsequent analyses, now referring to them simply as forests.

We used circular statistics that required converting time data into a radian scale, spanning from zero hours (0) to 24 hours (2π). To visualise diel activity patterns and subsequently calculate the overlap coefficients, we employed Kernel density functions. To do so, we used a smoothing parameter for Kernel density estimation (i.e., kmax) with a value of “three” (Ridout & Linkie, 2009, Rivero-Monteagudo & Mena, 2023). This value is known for robust estimates in both unimodal and bimodal activity distributions. We further applied an adjust factor of one for bandwidth scalar adjustment. These parameters interact to determine the width of the Kernel used for density estimation and later pattern display. The careful choice of *kmax* and adjust is crucial to accurately represent underlying patterns while avoiding undue influence from noise or over smoothing (Rivero-Monteagudo & Mena, 2023).

We then estimated overlap coefficients (Δ) that quantified (1) overlap between intraspecific activity between forest and human altered land-use types, and (2) overlap between interspecific species activity within the same land-use type. We considered three options of overlap coefficients, namely Dhat1 (Δ1), Dhat4 (Δ4), and Dhat5 (Δ5), each with distinct overlap measurement methods. Following Ridout and Linkie (2009), we selected Dhat4 due to more consistent performance across sample sizes. This overlap coefficient ranged from 0, meaning no overlap in diel species activity, to 1, indicating complete overlap. The confidence intervals with 95% certainty were estimated using the bootstrap technique with 9,999 resamples (Ridout & Linkie 2009). For these calculations, we used the *overlap* R package (Meredith & Ridout, 2014). To ascertain whether the two activity sets under comparison originated from identical distributions, we conducted a probability test using the ‘compareCkern’ function from the *Activity* package (Rowcliffe et al., 2014). All analyses were performed in R 4.1.2 software (R Core Team, 2022).

## 3. RESULTS

We recorded a total of 17,527 bat-passes from four species: the endemic edge-forager *Pseudoromicia principis* was the most detected species (15,165 bat-passes, 86.42%), followed by the open-space forager *Mops pumilus* (2,307 bat-passes, 13.16%), the forest forager *Hipposideros ruber* (48 detections, 0.27%), while *Taphozous mauritianus* was the least detected (7 detections, 0.04%). The two species *Pseudoromicia principis* and *Mops pumilus* were present in the four land-use types surveyed, while *H. ruber* was restricted to either old-growth or secondary re-growth forest (Fig. 2a). As *T. mauritianus* was only detected in three sites within shaded plantations (Fig. 2a), this species was not included in subsequent analyses.

**Figure 2.**
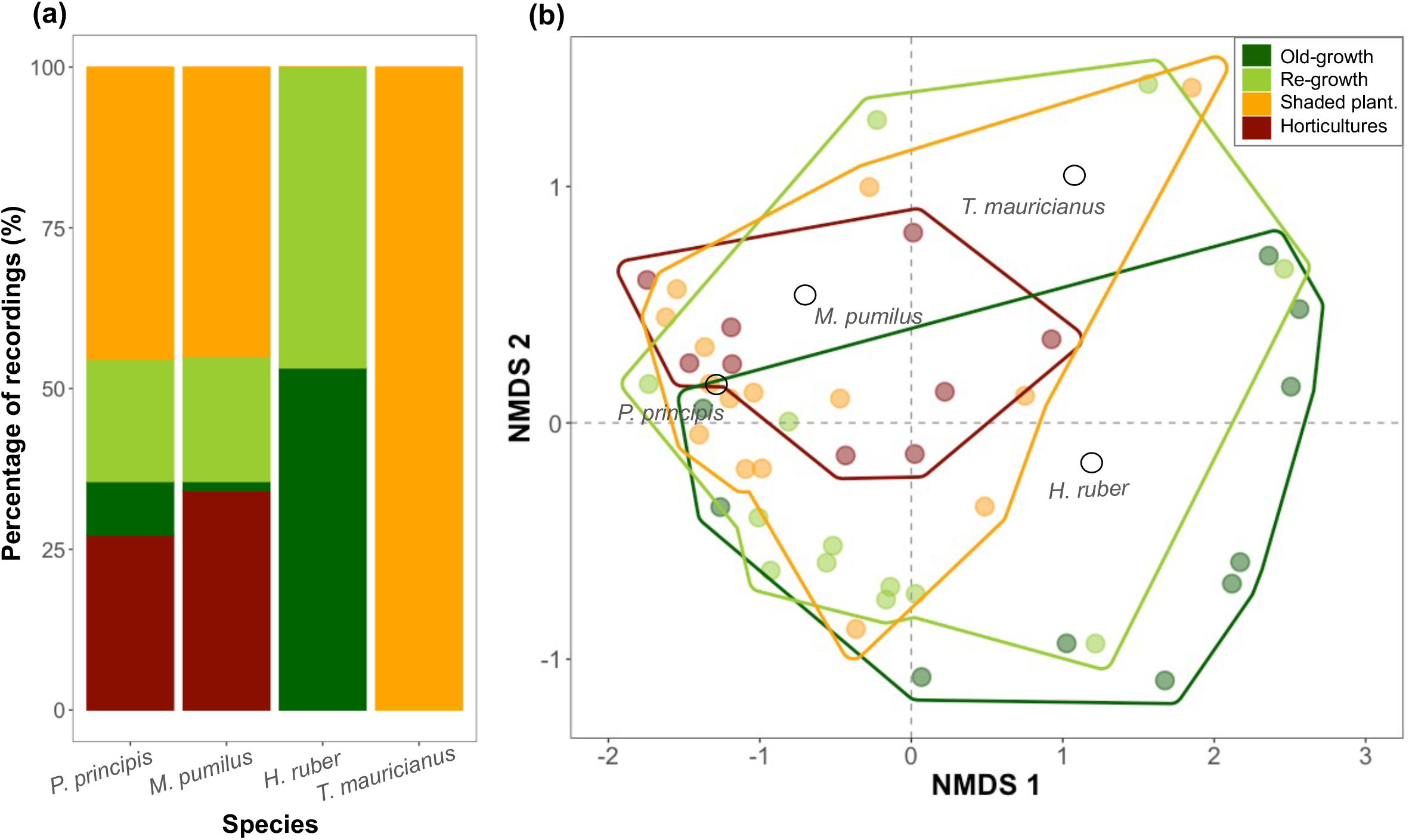
(a) Proportion (%) of the number of records for each of the four insectivorous bat species (i.e., *Pseudoromicia principis, Mops pumilus, Hipposideros ruber and Taphozous mauritianus*) recorded across the 48 sampling sites surveyed across all four major land-use types in Príncipe: old-growth forest, re-growth forest, cocoa shaded plantations, and horticultures. (b) Non-Metric Multi-Dimensional Scaling (NMDS) ordination plot denoting both sampling sites and species. In (b), sampling sites are represented by circles, color-coded according to land-use type.

### 3.1 Habitat use

Species composition was similar across land-use types (Fig. 2b, Table S2). Despite the apparent lower variation in species composition in horticultures, no differences in the species composition dispersion were observed (Fig. 2b, Table S2). Overall insectivorous bat activity was lower in old-growth forests in comparison to any of the remaining land-use types (Fig. 3a; Table S3), with no differences observed between these land-use types (Table S4)—a trend also observed for the most detected species, *P. principis* (Fig. 3b; Tables S3-S4). The activity of the open-space forager *M. pumilus* was lowest in both old-growth and re-growth forests in comparison to either shaded plantations or horticultures (Fig. 3c; Tables S3-S4). Lastly, *H. ruber* was similarly active in both land-use types where it was recorded, namely: old-growth and secondary re-growth forests (Fig. 3d; Tables S3-S4). Altitude had not effect on both overall activity nor species-specific bat activity (Table S3).

**Figure 3.**
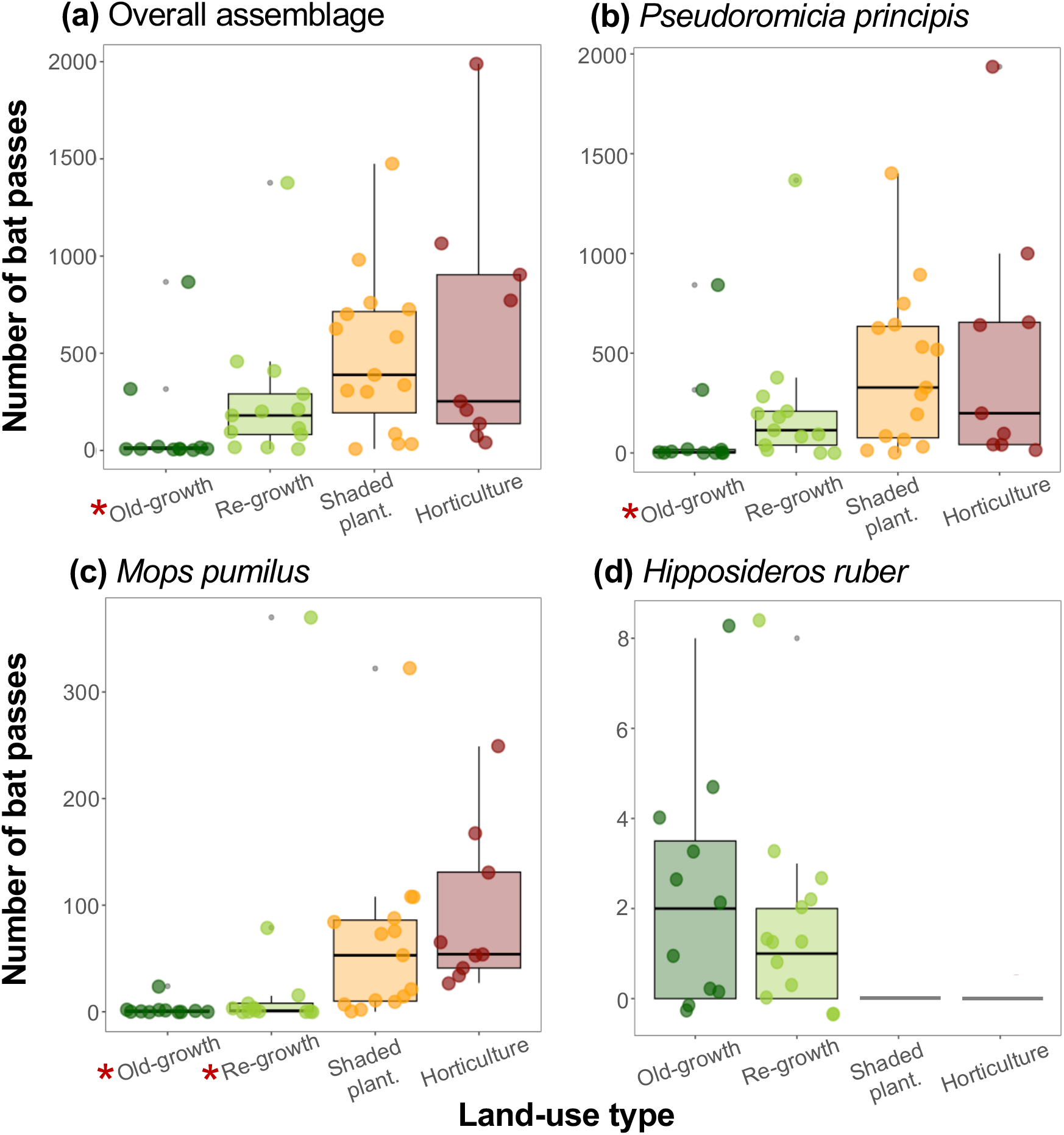
(a) Overall bat activity and species-specific activity of (b) *Pseudoromicia principis*, (c) *Mops pumilus*, and (d) *Hipposideros ruber* as denoted by the number of bat passes recorded across each of the four major land-use types in Príncipe: old-growth forest, re-growth forest, cocoa shaded plantations, and horticultures. Red asterisks indicate land-use types with significant lower number of bat passes in relation to the remaining land-use types.

### 3.2 Activity patterns

Species-specific activity patterns varied between forest and human-altered land-use types differently for *P. principis* and *M. pumilus* (Fig. 4). Indeed, the overlap in the activity of *P. principis* between forests and shaded plantations was as high as 0.94 (0.92 – 0.96) (Fig. 4a) and 0.90 (0.88 – 0.91) between forests and horticultures (Fig. 4b). On the contrary, the overlap in the activity of *M. pumilus* decreased from 0.80 (0.76 – 0.85) between forests and shaded plantations (Fig. 4c) to 0.44 (0.40 – 0.48) when considering forests and horticultures. In horticultures, *M. pumilus* presented two sharp activity peaks at dusk and dawn (Fig. 4d). Diel activity tended to be more uniform in forests. Forest had the highest overlap in activity across species with an overlap of 0.84 (0.80 – 0.88) (Fig. 5a), which decreased to 0.73 (0.70 – 0.76) (Fig. 5b) in shaded plantations, and to 0.47 (0.45– 0.50) in horticultures (Fig. 5c).

**Figure 4.**
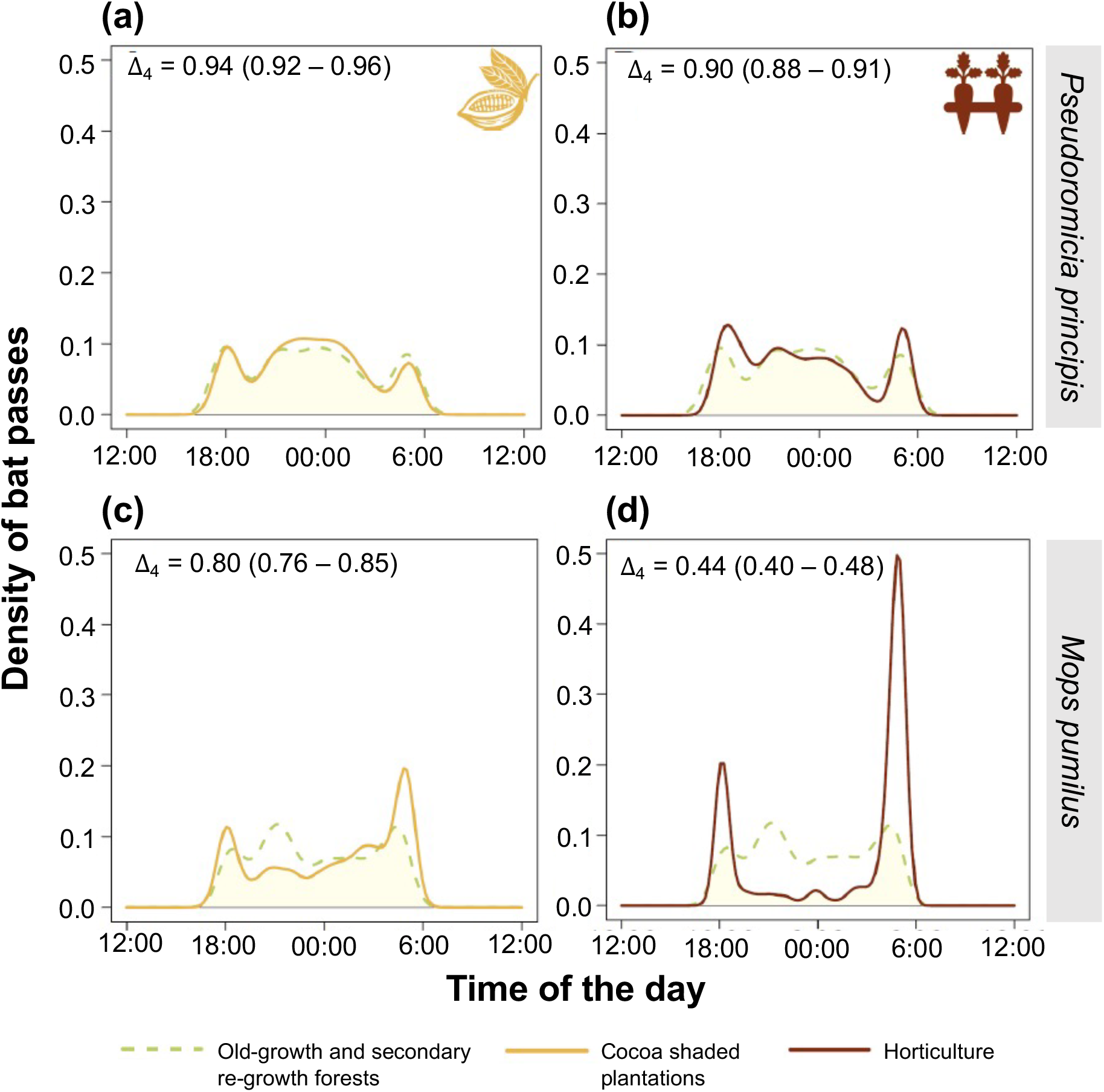
Comparison of the diel activity patterns of (a–b) *Pseudoromicia principis*, (c– d) *Mops pumilus*, between (a, c) forests (including old-growth and secondary re-growth forests) and cocoa shaded plantations, and between (b, d) forests and horticultures. For each comparison, the overlap coefficient and its confidence intervals are provided. All *P*-value testing the probability that the two or more distributions deriving from the same distribution were bellow 0.001.

**Figure 5.**
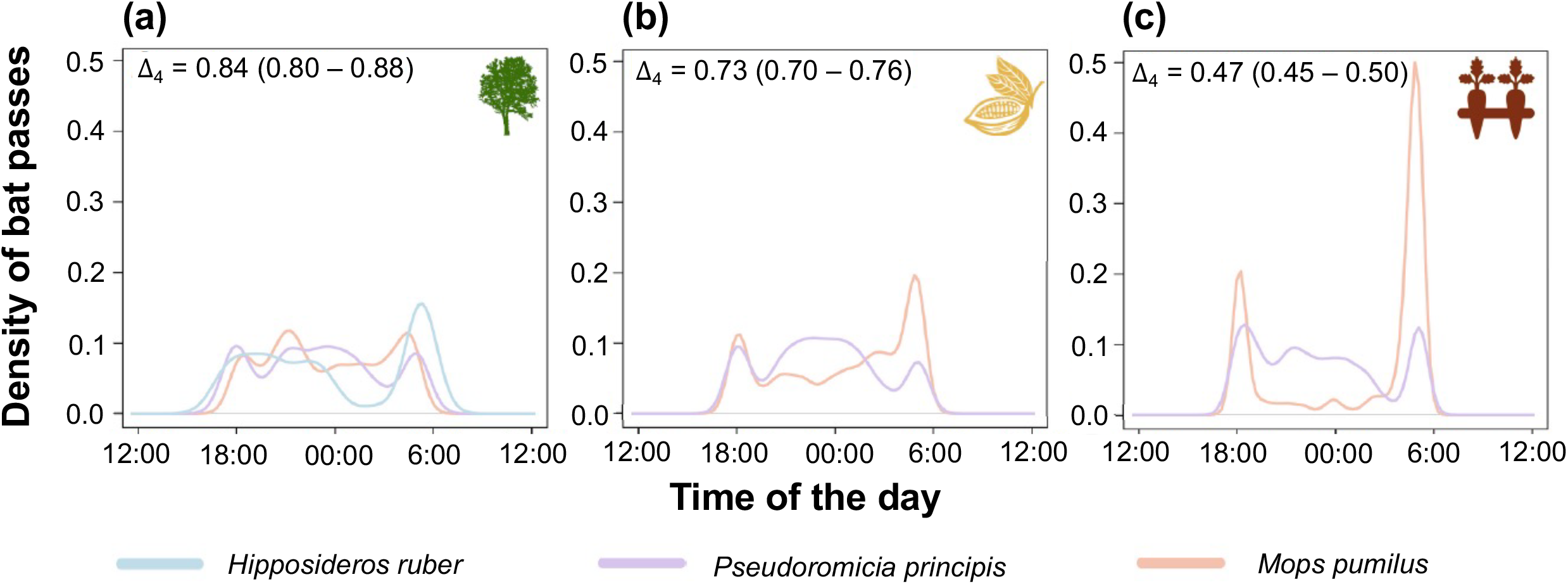
Comparison of the diel activity patterns of the species co-existing in each land-use type, including (a) forests (including both old-growth and secondary re-growth forests), (b) cocoa shaded plantations, and (c) horticultures. For each comparison, the overlap coefficient and its confidence intervals are provided. All *P*-value testing the probability that the two or more distributions deriving from the same distribution were bellow 0.001. Due to the low number of records, *Hipposiderus ruber* diel activity is presented in (a) but it was excluded from the analysis.

## 4. DISCUSSION

The exceptionally high biodiversity on islands, including a disproportionate number of threatened species (Ricketts et al., 2005), makes understanding the effects of land-use change in insular ecosystems critical to adverting the ongoing defaunation crisis (Fernández-Palacios et al., 2021). The current lack of knowledge is particularly acute on Afrotropical islands (de Lima et al., 2011; Conenna et al., 2017), many of which—such as the ones located in the Gulf of Guinea—host exceptional levels of endemicity. Here, we detected two of Príncipe’s bat species in each of the major land-use types. While for three of the four species had low or no activity in old-growth forest, the forest forager *H. ruber* was exclusively found in forested habitats, highlighting the importance in habitat diversity for different species in the bat community. The endemic *P. principis* showed little difference in diel activity patterns across habitat types, suggesting is it able to exploit resource opportunities across human-modified landscapes as found in forested habitats. Diel activity patterns of *M. pumilus* varied between forests and non-forest land-use types, whereas those of *P. principis* remained similar. Interspecific activity overlaps further decreased towards more altered land-use types.

### 3.1 Habitat use

Overall insectivorous bat activity was lowest in old-growth forests. In fact, despite the exclusivity of *H. ruber* to forests, the two more abundant species, *P. principis* and *M. pumilus* were mainly recorded in cocoa shaded plantations and horticultures. Both species are well-adapted to exploit resources in more open-spaces, but less able to forage in structural complex vegetation characteristic of old-growth forest. While several studies have reported negative effects of habitat modification on island bats (e.g., Davies et al., 2016; Ferreira et al., 2022; Moseley et al., 2022), our findings are consistent with those observed for insectivorous bat assemblages in the nearby São Tomé Island, where overall activity was higher the same non-forest land-use types (Fernandes et al., under review). Likewise, a higher species richness and abundance of (non-endemic) birds (de Lima et al., 2013; Soares et al., 2020), as well as a higher abundance of amphibians (Strauß et al., 2018) was also reported for the anthropized habitats of São Tomé and elsewhere across the tropics (e.g., Silva-Souza et al., 2022; Kemp. et al., 2019). Therefore, human modification may increase opportunities for certain species. For example, secondary re-growth forests are broadly characterised by the presence of fruiting trees, which might boost overall insect availability and increase prey for *P. principis* and *M. pumilus* (Duque-Trujillo et al., 2022). Further research is needed to assess activity in relation to prey availability across these landscapes.

As already observed on São Tomé Island (Fernandes et al., under review), shaded plantations and horticultures exhibited the highest activity of insectivorous bats. Shaded plantations, resembling agroforests, involve cocoa cultivation beneath the canopy of predominantly exotic trees (de Lima et al. 2013). Horticultures consist mainly of smallholding fields, typically forming heterogeneous mosaics of diverse crops such as banana, pineapple, and cassava. Both habitat types are characterized by low-intensity management, with minimal to no use of pesticides or fertilizers. Organic management practices are known to benefit (pollinator) insects (Duque-Trujillo et al., 2023), eventually resulting in a higher trophic resource availability for insectivorous bats (Williams-Guillén et al., 2016). The molossid *M. pumilus* are particularly well adapted to fly in open areas due to the high aspect ratio (long and narrow) of their wings (Jung and Kalko, 2011) and as a result broadly benefit from some less structurally complex habitat types (Russo and Ancillotto, 2015). In fact, molossids often tend to comprise the most recorded bats in horticultures (Kemp et al., 2019). This species adaptability to humanized habitats might further relate to their roost plasticity ranging from house roofs to tree cavities, palm leaves and rocky crevices (Kingdon 1974). *M. pumilus* is known to play an important role as suppressor of agricultural pests (Wanger et al., 2014), thus likely benefiting of an increased abundance of pest arthropods in the island’s agricultural matrix.

The Príncipe endemic *P. principis* is known to be relatively abundant in forest, horticultures and urban areas (Rainho et al., 2022). This small vespertilionid is an edge forager—a guild that is typically flexible in their foraging and echolocation behaviour, frequently switching between open and clutter-edge habitats (Fenton, 1990). It is likely that these characteristics enable *P. principis* to thrive, not only in along forest edges, but also in human-altered land-use types, thereby explaining its island-wide distribution.

The open-space forager *T. mauritianus* was only recorded seven times in three sites located within shaded plantations. Despite being relatively rare on Príncipe, this species is not known to be highly sensitive to land-use change. Noteworthy, although an efficient island colonizer (Bonaccorso, 2019), the presence of *T. mauritianus* was only recently confirmed in Príncipe (Rainho et al., 2022).

The forest forager *H. ruber* was restricted to both old-growth and secondary re-growth forests (but see Rainho et al., 2022). Forest specialists, such *H. ruber*, tend to have short, broad wings and emit high-frequency, broadband echolocation calls, which attenuate quickly but offer more detailed information—traits that are particularly well-suited for navigating and hunting in cluttered environments (Denzinger and Schnitzler, 2013; Mande et al., 2023). These specializations, along with additional life-history traits (e.g. reliance on tree roosts), mean these species are unable to persist in non-forest habitats. Therefore, it is vital that forests are preserved within human-modified landscapes.

An important caveat to our findings is that bat calls are harder to detect in cluttered environments (Duchamp et al., 2006), which might have also contributed to the lower overall bat activity recorded in both old-growth and secondary re-growth forests compared to more open habitats. Similarly, higher frequency calls attenuate more quickly than lower frequency calls, therefore reducing the detectability of forest specialist species two-fold (Russo et al., 2018). Another consideration is vertical stratification exhibited across species. *M. pumilus* forages above the tree canopy (Duchamp et al., 2006), which might have impacted the species’ detection in forest sites as we deployed detectors in the understory.

### 4.2 Diel activity patterns

While *P. principis* maintained similar uniform diel activity patterns across forests and non-forest habitats, *M. pumilus* differed from uniform diel activity in forests to a diel activity characterised by dusk and dawn peaks in non-forested habitats, most pronounced in horticultures. As the open-space forager *M. pumilus* is poorly suited to forest habitats, it is possible that the diel activity pattern observed in the horticultures corresponds to the usual pattern of this species, generally driven by canopy cover, light, (Russo et al., 2007) and arthropod diversity/activity (Presley et al., 2009, Montaño-Centellas et al., 2015). In addition, interspecific competition tends to intensify at higher species abundances, in which case species might be forced to displace their temporal habitat use (Lambert et al., 2018). This might explain the fact that *P. principis* is often more active in periods when *M. pumilus* is less so in shaded plantations and horticultures.

Similarly, elsewhere in the tropics, and including in the nearby island São Tomé, interspecific diel activity overlap of insectivorous bats was also lower in more species-rich habitats, with greater forest cover (Rivero-Monteagudo & Mena 2023; Araújo-Fernandes et al., under review).

Furthermore, we did not observe any diurnal activity of *H. ruber* on Príncipe during our study period. This contrasts with findings from São Tomé, where the species is largely active during both the day and night (Russo et al., 2011; Rainho et al., 2022; Araújo-Fernandes et al., under review). Both islands lack specialized diurnal avian bat predators, thus this disparity in behaviour is rather intriguing and requires further study as it might signal more significant population-level differences.

### 4.3 Implications for conservation

Forest conversion into working landscapes is a major driver of the contemporary biodiversity crisis (Caro et al., 2022)— particularly for oceanic islands (Fernández-Palacios et al., 2021). On Príncipe, where only 25% of the native island forest cover persists (Dauby et al., 2022), the extant insectivorous bat fauna is persisting under the current land-use configuration. To sustain current bat activity levels within non-forest land-use types, we support the continued use of established low intensity management practices. In addition, given the particularly high activity of insectivorous bats in both shaded-plantations and small-holdings horticultures, we highlight the need to understand their potential role as pest suppressors (Kemp et al., 2019, Ferreira et al., 2023). Yet, as the forest forager *H. ruber* was restricted to the forests of Príncipe, our results also emphasize the role native habitats play to preserve island-wide integrity of bat assemblages. In summary, preserving a mosaic of land-use types, including native forests, would help optimize the long-term survival of bat species in Príncipe, a strategy that may also be effective on other human-populated, tropical oceanic islands.

## Acknowledgments

AFP, ASF, ACF, PG, and RR and were supported by the European Union’s Horizon 2020 research and innovation programme under grant agreement No 854248 (TROPIBIO). This programme further funded the fieldwork. VAM was supported by Fundação para a Ciência e a Tecnologia (FCT) through an individual research contract (https://doi.org/10.54499/2020.02547.CEECIND/CP1601/CP1649/CT0004). We thank the Direction of Forests and Biodiversity of São Tomé and Príncipe for authorising the surveys. Additionally, we thank all the landowners that allowed us to survey in their properties.

