## Supplementary material for "Habitat use and diel activity of insectivorous bats across land-use types on an Afrotropical oceanic island"

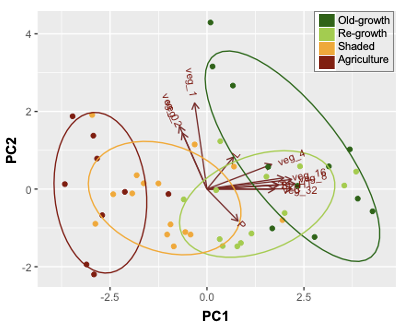

**Figure S1.** Diagram of the Principal Component Analysis (PCA) based on the characteristics of each of the 48 sampling sites located in old-growth forest, re-growth forest, cocoa shaded plantations and horticultures in Princípe Island, Central West Africa. Sampling sites are colour-coded according to the land-use type where the site was located. The strength of each habitat characteristic considered is represented by the red arrows and the name of the habitat characteristic is abbreviated (the full name of habitat characteristics can be found in Table S1). The axis 1 of the PCA (PC1) explained 38.2% of the variance and second axis (PC2) explained 13.3% of the variance.

**Table S1.** Summary of local-scale variables. T = number of trees (DBH ≥ 10 cm), HT = Tree height (m), S = number of woody stems (DBH < 10 cm), P = number of palms, C = percent canopy cover, L = number of lianas, VC = vegetation cover at 0, 1, 2, 4, 8, 16, and 32 m. Results are presented as mean ± SD.

| **Variable** | **Old-growth forest** | **Secondary re-growth forest** | **Cocoa shaded plantation** | **Horticulture** |
| --- | --- | --- | --- | --- |
| **T** | 11.67 ± 3.55 | 10.00 ± 7.79 | 6.33 ± 5.11 | 4.00 ± 4.14 |
| **HT** | 30.15 ± 7.47 | 20.23 ± 6.60 | 17.87 ± 5.66 | 9.98 ± 5.82 |
| **S** | 27.42 ± 12.09 | 29.87 ± 15.96 | 2.60 ± 4.31 | 3.70 ± 5.56 |
| **P** | 4.83 ± 5.46 | 10.40 ± 5.44 | 5.07 ± 6.02 | 1.60 ± 2.01 |
| **C** | 85.42 ± 7.53 | 80.00 ± 9.26 | 49.67 ± 25.03 | 22.50 ± 20.85 |
| **L** | 0.92 ± 0.67 | 0.73 ± 0.80 | 0.40 ± 0.91 | 0.2 ± 0.63 |
| **VC 0m** | 29.17 ± 24.01 | 16.67 ± 16.55 | 15.00 ± 10.52 | 40.50 ± 28.33 |
| **VC 1m** | 27.08 ± 17.38 | 20.67 ± 12.37 | 18.33 ± 14.84 | 20.00 ± 13.12 |
| **VC 2m** | 22.50 ± 13.06 | 19.00 ± 9.49 | 34.33 ± 18.89 | 19.50 ± 13.22 |
| **VC 4m** | 36.25 ± 14.64 | 34.33 ± 14.98 | 23.33 ± 16.55 | 16.00 ± 13.29 |
| **VC 8m** | 39.58 ± 10.97 | 40.67 ± 17.10 | 19.00 ± 15.02 | 10.50 ± 15.36 |
| **VC 16m** | 40.83 ± 16.76 | 25.33 ± 19.13 | 13.00 ± 15.56 | 1.50 ± 3.37 |
| **VC 32m** | 15.00 ± 13.65 | 1.67 ± 4.50 | 1.00 ± 3.87 | 0 |

**Table S2.** Results of the PERMANOVA and PERMDIST.

| **PERMANOVA** | | **Df** | **Sum Of Sqs** | **R2** | **F** | **Pr(>F)** |
| --- | --- | --- | --- | --- | --- | --- |
|  | Group vector | 3 | 1.3492 | 0.09475 | 1.5352 | 0.101 |
|  | Residual | 44 | 12.8902 | 0.90525 |  |  |
|  | Total | 47 | 14.2394 | 1 |  |  |
| **PERMDIST** |  | **Df** | **Sum Sq** | **Mean Sq** | **F** | **Pr(>F)** |
|  | Groups | 3 | 0.08153 | 0.027175 | 0.768 | 0.522 |
|  | Residuals | 44 | 1.55683 | 0.035383 |  |  |

**Table S3.** Results of Generalised Linear Models (GLMs) relating overall insectivorous bat activity and the activity of *Pseudoromicia principis*, *Mops pumilus* and *Hipposideros ruber* with land-use type, including old-growth forests, re-growth forests, cocoa shaded plantations, and horticultures in the Princípe Island. For each model, we indicate the estimate, standard error, z-value, and P-value.

| **Response** | **Model parameters** | **Estimate** | **Std. error** | ***z*-value** | ***P*-value** |
| --- | --- | --- | --- | --- | --- |
| *Overall activity* | |  |  |  |  |
|  | Intercept | 3.108 | 0.774 | 4.016 | <0.0001 |
|  | Re-growth | 1.844 | 0.678 | 2.719 | 0.007 |
|  | Shaded | 2.677 | 0.630 | 4.252 | <0.0001 |
|  | Horticulture | 2.949 | 0.669 | 4.408 | <0.0001 |
|  | Altitude | -0.002 | 0.003 | -0.501 | 0.617 |
| *Pseudoromicia principis* | |  |  |  |  |
|  | Intercept | 1.843 | 0.728 | 2.533 | 0.011 |
|  | Re-growth | 2.145 | 1.020 | 2.104 | 0.035 |
|  | Shaded | 3.295 | 0.937 | 3.515 | 0.000 |
|  | Horticulture | 3.532 | 0.987 | 3.579 | 0.000 |
|  | Altitude | -0.242 | 0.366 | -0.660 | 0.509 |
| *Mops pumilus* | |  |  |  |  |
|  | Intercept | -0.565 | 0.723 | -0.781 | 0.435 |
|  | Re-growth | 1.117 | 0.934 | 1.196 | 0.232 |
|  | Shaded | 3.726 | 0.869 | 4.286 | 0.000 |
|  | Horticulture | 5.028 | 0.922 | 5.453 | 0.000 |
|  | Altitude | -0.495 | 0.328 | -1.512 | 0.130 |
| *Hipposideros ruber* | |  |  |  |  |
|  | Intercept | 0.841 | 0.404 | 2.083 | 0.037 |
|  | Re-growth | -0.299 | 0.615 | -0.486 | 0.627 |
|  | Altitude | 0.029 | 0.313 | 0.093 | 0.926 |

**Table S4.** Results of the comparisons of insectivorous bat activity between pairs of land-use types. Comparisons were made based on the Generalised Linear Models summarised in Table S3. Comparisons are presented for overall insectivorous bat activity and that of *Pseudoromicia principis* and *Mops pumilus.* For each comparison, we indicate the contrast between the estimates, standard error, degrees of freedom, t-value, and P-value.

| **Response** | **Comparison** | **Constrast** | | **SE** | | **df** | | ***t*-value** | | ***P*-value** |
| --- | --- | --- | --- | --- | --- | --- | --- | --- | --- | --- |
| *Overall activity* | | |  | |  | |  | |  | |
|  | Re-growth x Shaded | -0.833 | | 0.937 | | 42 | | -0.889 | | 0.379 |
|  | Re-growth x horticulture | -1.105 | | 0.963 | | 42 | | -1.148 | | 0.258 |
|  | Shaded x horticulture | -0.272 | | 0.927 | | 42 | | -0.293 | | 0.771 |
| *Pseudoromicia principis* | | |  | |  | |  | |  | |
|  | Re-growth x Shaded | -1.150 | | 1.385 | | 42 | | -0.830 | | 0.411 |
|  | Re-growth x horticulture | -1.386 | | 1.419 | | 42 | | -0.977 | | 0.334 |
|  | Shaded x horticulture | -0.237 | | 1.361 | | 42 | | -0.174 | | 0.863 |
| *Mops pumilus* | | |  | |  | |  | |  | |
|  | Re-growth x Shaded | -2.610 | | 1.276 | | 42 | | -2.046 | | 0.047 |
|  | Re-growth x horticulture | -3.911 | | 1.312 | | 42 | | -2.981 | | 0.005 |
|  | Shaded x horticulture | -1.301 | | 1.267 | | 42 | | -1.027 | | 0.310 |
